# How strategy environment and wealth shape altruistic behaviour: Cooperation rules affecting wealth distribution in dynamic networks

**DOI:** 10.1101/2020.05.05.077131

**Authors:** Spandan Pathak, Prateek Verma, Sumit K. Ram, Supratim Sengupta

## Abstract

Societies rely on individual contributions to sustain public goods that benefit the entire community. Several mechanisms, that specify how individuals change their decisions based on past experiences, have been proposed to explain how altruists are not outcompeted by selfish counterparts. A key aspect of such strategy updates involves a comparison of an individual’s latest payoff with that of a random neighbour. In reality, both the economic and social milieu often shapes cooperative behaviour. We propose a new decision heuristic, where the propensity of an individual to cooperate depends on the local strategy environment in which she is embedded as well as her wealth relative to that of her neighbours. Our decision-making model allows cooperation to be sustained and also explains the results of recent experiments on social dilemmas in dynamic networks. Final cooperation levels depend only on the extent to which the strategy environment influences altruistic behaviour but are largely unaffected by network restructuring. However, the extent of wealth inequality in the community is affected by a subtle interplay between the environmental influence on a person’s decision to contribute and the likelihood of reshaping social ties, with wealth-inequality levels rising with increasing likelihood of network restructuring in some situations.

## 1. Background

Contemporary society faces many challenges like climate change, pandemics and mass migration due to social conflicts that require concerted action not only from governments but also from individuals. Growing wealth inequality is another pressing problem that can undermine democratic norms leading to destabilization of society by increasing the number and intensity of social conflicts. A recent poll by Ipsos[1] reveals that the wealthiest 1% own more than 30% of the total household wealth in 17 of 29 countries surveyed with disparities being even greater in many major emerging economies. Individuals as well as nation states are often faced with the choice of contributing to the public good for the benefit of society. Such disparities in wealth can affect the quantum of contribution necessary for sustaining various public goods thereby lowering the quality of life and the environment[2]. Hence, our actions as individuals as well as those of our government can potentially affect our ability to survive[3]. In view of their significance, it is important to understand how the factors that affect individual choices shape collective outcomes. Here, we specifically explore how sustenance of cooperation and wealth inequality levels in the population are affected when a person’s altruistic behaviour is informed by the strategies of connected neighbours and the wealth of the neighbourhood.

Various mechanisms like direct reciprocity[4,5], indirect reciprocity[6–8], kin selection[9] and structured populations[10–13], have been put forward to explain the evolution of cooperation[14]. All these mechanisms specify rules for updating strategies in response to past experiences of interactions and in some cases, the reputation of interacting partners[7,15]. However, the effectiveness of these update rules in sustaining cooperative behaviour can depend on the structure of the underlying social network as well. Several studies have highlighted the usefulness of rewiring social ties[16–25] in promoting cooperation in the absence of punishment. While these papers differ in the details of the strategy update and rewiring processes, they all show that the fraction of cooperators in the population can be enhanced by the coevolution of individual strategy and the underlying network structure. In all these models, strategy update occurs after a pair-wise comparison of payoffs between the focal player and a randomly chosen neighbour. Low-information, learning-based[26] models, where strategy-update rules depend on individual satisfaction levels[27,28] that are oblivious to the payoffs or reputation of connected neighbours, have also been proposed. They reveal that cooperation and social cohesion can be sustained even at moderate levels of greediness despite lack of information about strategies of connected neighbours.

The need to account for social diversity[29], resource [30] and wealth [2,31–33] heterogeneity in understanding altruistic behaviour, is increasingly being recognized. Such differences in wealth, and benefits received from public resources on the evolution of altruistic behaviour can have widespread consequences. High wealth inequality leads to more exploitation of poorer people by richer people while the rich tend to be more generous when wealth inequality is low[34]. Removing the anonymity of wealth accumulated by players can lead to increased donations by richer participants[35]. On the contrary, the visibility of the wealth of connected neighbours was found to adversely affect equilibrium wealth distribution and cooperation levels in co-evolving social networks[36]. An aspect of strategic decision-making that has received far less attention is the extent to which the evolving strategy environment of a player affects her altruistic behaviour. Nevertheless, an influential study[37] showed that the presence of unconditional cooperators in a networked PGG was found to systematically increase contributions of other players, thereby pointing to the existence of conditional cooperators who’s behaviour is influenced by cooperators in the neighbourhood. Even though this study did not examine the effect of changes in the number of altruistic neighbours, it provides circumstantial evidence to support the importance of strategy environment on altruistic behaviour. Behavioural experiments[38–40] have also pointed out striking disparities between the “imitate the best” predictions of strategy update based on the pairwise comparison rule[41,42] and the actual behaviour of human participants playing prisoners’ dilemma games. These studies underscore the need to come up with decision heuristics that more accurately reflects the evolution of human behaviour in social dilemmas. They enabled us to identify three key factors (wealth, local strategy environment and network restructuring) that need to be incorporated into any decision-making model that attempts to understand how cooperation can be sustained.

In this paper, we use a variant of a public goods game (PGG) to present a new model for cooperative strategy updating that depends on the relative wealth and the nature of the local strategy environment in which the individual is embedded. The local strategy environment is a cooperative one if ≥50% of neighbours are cooperators but it is a selfish one >50% of neighbours are defectors (see Fig.1). Our model provides a striking and testable alternative to the decision-making framework based on pairwise-comparison of payoffs, that is typically used in evolutionary game theory models (see the next section). We find that such a decision heuristic can sustain cooperation with cooperation levels being determined by the impact of the local environment on individual decisions. An increasing likelihood of reorganizing social ties does not affect cooperation levels but does affect final wealth-inequality in a manner that depends on the role of the strategic environment on altruistic behaviour. Fig. 1 highlights the key features of our model (see also supplementary figure S1). Supplementary Video1 shows the evolution of individual strategy and wealth in a dynamic network.

**Figure 1.**
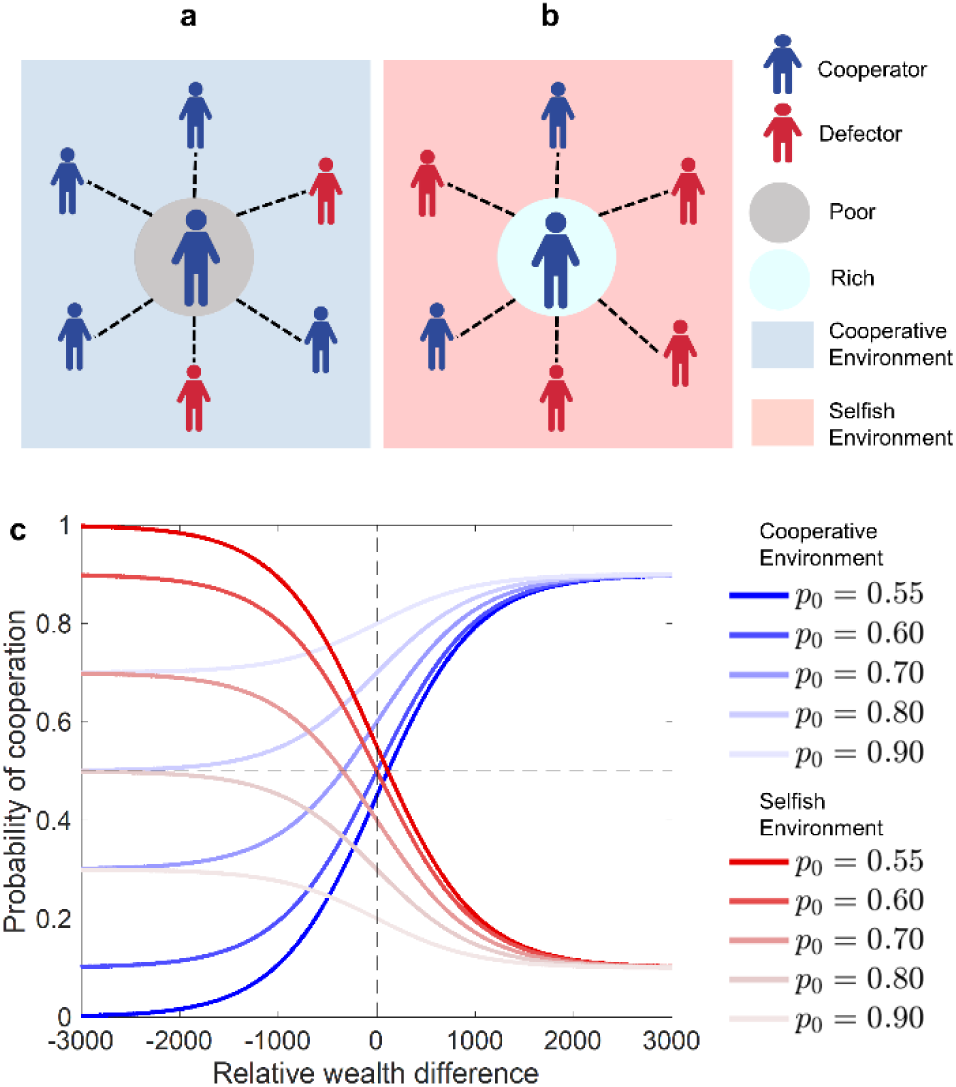
Salient features of the model: (a) A poor cooperator in a cooperative environment and (b) A rich cooperator in a selfish environment. (c) The S-shaped function showing the impact of the wealth difference (relative to the average wealth of the neighbourhood) and the strategy environment (quantified by p_0_) on the likelihood of cooperation.

## 2. Methods

Our model consists of individuals embedded as nodes in a random (Erdos-Renyi) network playing a PGG with each connected neighbour over multiple rounds. The model, which closely follows the experimental protocol described in [36], can be constructed in 3 stages : (i) Initial network formation and wealth allocation. (ii) Decision step in which each player with *k* connections plays (*k+1*) PGG games, once as a focal player and the remaining *k* games as a neighbour of a focal player (iii) Network rewiring step where a specified fraction of pairs of individuals are given the option to restructure the network through making or breaking links. A pictorial representation of the model along with additional details can be found in the electronic supplementary material (Figure S1).

The Initial network structure is created by imposing the condition that any two randomly selected members are connected with a probability *q=0*.*3*. After network formation, each member is initially assigned one of two possible wealth levels (rich and poor) and provided with an amount of money (endowment) to match the initial wealth distribution in the population. The level of wealth inequality in the population is measured by the Gini coefficient 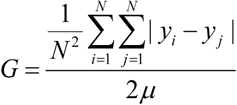 with the initial distribution fixed by specifying an *initial* Gini coefficient. *y*_*i*_ is the wealth of the i’th individual and m is the mean of the wealth distribution. G=0 implies perfect equality with every member having the same wealth and *G=1* corresponds to perfect inequality with one member possessing the entire wealth.

### Decision step

In the decision step, following [36], each individual plays a modified public-goods game (PGG) with all of her neighbours. During each such game, the individual can choose to either cooperate (C) by pledging a fixed amount of money (m=50 units) to each connected neighbour or defect (D) by refusing to donate. The amount pledged is multiplied by a synergy factor (*r*) and redistributed to all the connected neighbours regardless of their choice to cooperate or defect as focal players. If the focal individual chooses to defect, none of her connected neighbours get anything. The payoff to the i’th individual (C or D), connected to *n*_*c*_ cooperators and (*k*_*i*_*-n*_*c*_) defectors, is given by 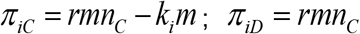. In the first round, every individual has a fixed probability (*p*_*C*_*=0*.*7*) of cooperating which reflects a collection of people who choose to be mostly cooperative initially on the basis of trust, although cultural differences may affect the way in which different people behave[43]. In subsequent rounds, individuals take cues from their strategy environment while deciding whether to cooperate. The cooperation probability of a focal player depends on the fraction of C neighbours and on the comparison between her wealth and the average wealth of her connected neighbours. On rare occasions, a focal player may have equal number of C and D neighbours, in which case the strategy environment is defined as a neutral one. Previous work[34] suggests that rich and poor people behave differently with richer people being less generous especially in high wealth-inequality scenarios. A poorer player (relative to the average wealth of the local neighbourhood) has a higher incentive to cooperate with her neighbours since doing so increases the likelihood of her receiving favours from them in return. On the other hand, a richer player has a higher incentive to exploit her neighbours by defecting, since her higher wealth ensures that she pays a relatively lower cost when a link with her is broken in response to her selfish behaviour. Hence a poorer player is influenced by the wisdom of the majority and is likely to cooperate with a higher probability (*p*_*C*_*>0*.*5*) if her local environment is mostly cooperative (≥50% C neighbours) and more likely to defect if her local environment is mostly selfish (<50% C neighbours). By contrast, a richer player has more to gain by defecting in a cooperative environment and a greater likelihood of cooperating in a predominantly selfish environment to reduce the chances of being isolated due to breaking of her existing links. We therefore propose the following stochastic decision-making rule that gives the probability of cooperation (*p*_*C*_) for a focal player that is dependent on her local strategy environment and relative wealth difference.

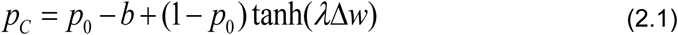

when the focal player is in a cooperative environment and

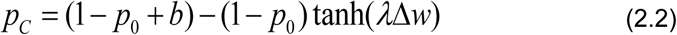

when the focal player is in a selfish environment. Δ *w* = *E*_*LC*_ (*w*) - *w*_*F*_ is the relative wealth difference; *E*_*LC*_ (*w*) is the average cumulative wealth of neighbours of the focal player and *w*_*F*_is the focal player’s cumulative wealth accumulated over past rounds. *p*_*0*_ determines the influence of the strategy environment on an individual’s propensity to cooperate (see Fig.1c) and *l* determines the extent to which the wealth of the local environment affects the focal player’s likelihood of cooperation. *l*=0 implies that the focal player’s decision is insensitive to the wealth of the local neighbourhood. *b* is a bias term that regulates a player’s maximum and minimum propensities for cooperating. *b≠0* ensures that a considerably poorer player, whose wealth is very small compared to that of her neighbours, does not always cooperate in a cooperative environment.

Our decision rule (Eq. 2.1 & 2.2) is to be contrasted with the pairwise comparison rule. In the latter case, the decision to cooperate is made after a comparison of payoffs with a randomly chosen neighbour with the probability of changing strategies given by 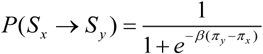 where P(*S*_*x*_→*S*_*y*_) is the probability that the focal player x adopts the strategy of a neighbour y; π_x_, π_y_ are the respective payoffs and β is a measure of the intensity of selection. The local environment plays no role, contrary to our case where the local strategy environment influences cooperative behaviour.

### Network rewiring step

Following the decision update step, a fraction *re* (henceforth called the rewiring fraction) of all possible pairs of individuals chosen randomly are given the *option* to rewire their connections. If the pair of selected nodes is already connected via a link, then one of them is selected at random to take the decision about retaining or breaking the link. If the partner of the randomly chosen decision-maker had cooperated in the previous round, the link is retained with probability *p*_*r*_, otherwise the link is broken with probability *p*_*b*_. If the selected pair of nodes are not connected by a link, both are given a choice to establish a new connection. A link is established with a high probability *p*_*m*_ if *both* had cooperated in the previous round. If either or both of the nodes had defected in the last round, a new link can still be established with probability *p*_*e*_ and *p*_*s*_ respectively. Unless specified otherwise, we used *p*_*r*_=0.87, *p*_*b*_=0.7, *p*_*m*_=0.93, *p*_*e*_=0.3, *p*_*s*_=0.2 for the network restructuring probabilities. Changes in these values within reasonable bounds do not affect our conclusions.

## 3. Results

Fig. 2 highlights how our decision rule, which relies on comparison between the focal player’s wealth and the average wealth in her local network neighbourhood, leads to distinctly different outcomes compared to the standard pairwise comparison rule[41,42] used in evolutionary game theory. Our proposed decision heuristic ensures that the fraction of cooperators stabilizes to around 0.59 in contrast to the pairwise comparison rule modelled using a Fermi function where the fraction quickly decays to zero. This fraction depends on the minimum and maximum cooperation levels for relatively richer and poorer individuals and is controlled by the environmental parameters *p*_*0*_ and *b*. Wealth inequality also reduces in our model in contrast to the standard decision-update rule where it saturates to a higher value. A change in the initial Gini coefficient does not change the qualitative behaviour in these cases. A greater reduction in wealth inequality observed in our model can primarily be attributed to the higher fraction of cooperators and a large number of local cooperative environments at equilibrium, as well as an increase in the average degree of the network in comparison to the standard case. These three features ensure more efficient accumulation and redistribution of wealth generated from each PGG, consequently leading to lower wealth-inequality.

**Figure 2.**
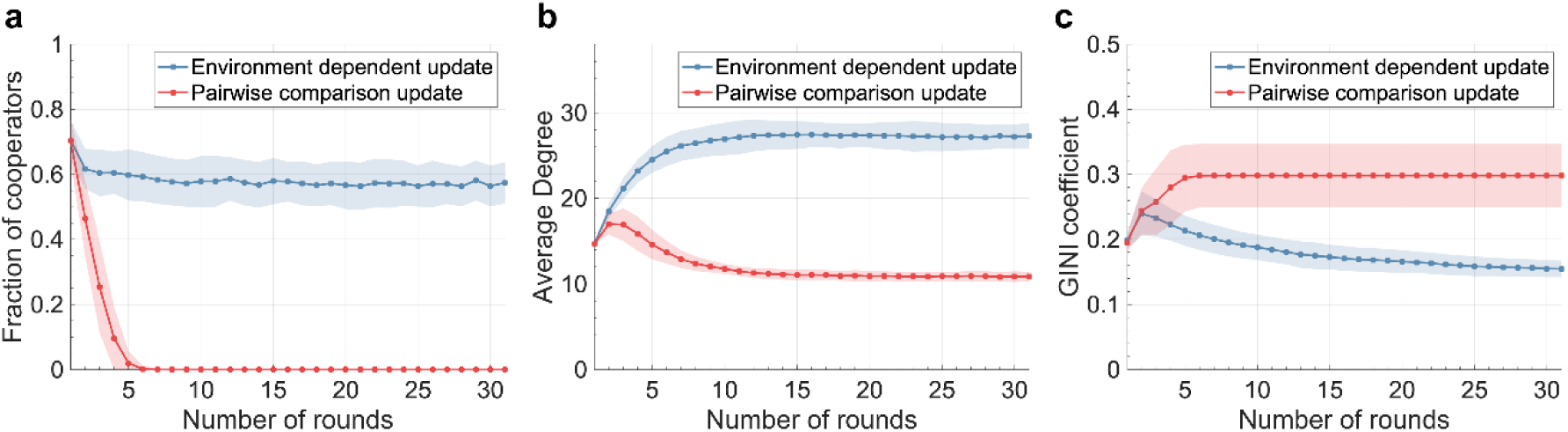
Comparison between the environment and wealth-dependent decision heuristic and the pairwise comparison rule: (a) The fraction of cooperators and (b) the average degree saturate to a comparatively higher value while (c) the wealth inequality reduces significantly, in the former case. These results are independent of the initial wealth-inequality levels and have been obtained by averaging over 100 trials. The shaded regions represent one sigma variation from the mean. Parameters used: N=50, β=0.01, b=0.1, p_0_=0.7, λ=0.001, initial Gini=0.2, re=0.3, r=2.

Fig. 3a,b shows heat maps depicting the equilibrated fraction of cooperators and extent of wealth inequality in the population (quantified by the final Gini) when the influence of the local strategy environment and the rewiring fraction are varied. Once the impact of the strategy environment on decision making is fixed (by specifying p_0_), an increase in the probability of changing network structure by rewiring of social ties does not have much of an impact on the frequency of cooperators. However, the extent to which wealth-inequality is affected by changing this rewiring fraction depends on the impact of the strategy environment. As we explain in the next two sub-sections, these results can be understood by carefully analyzing how the strategy update dynamics affects network topology manifest through the connectedness of rich and poor cooperators and defectors as well as the nature (C or D) of the dominant strategy environments.

**Figure 3:**
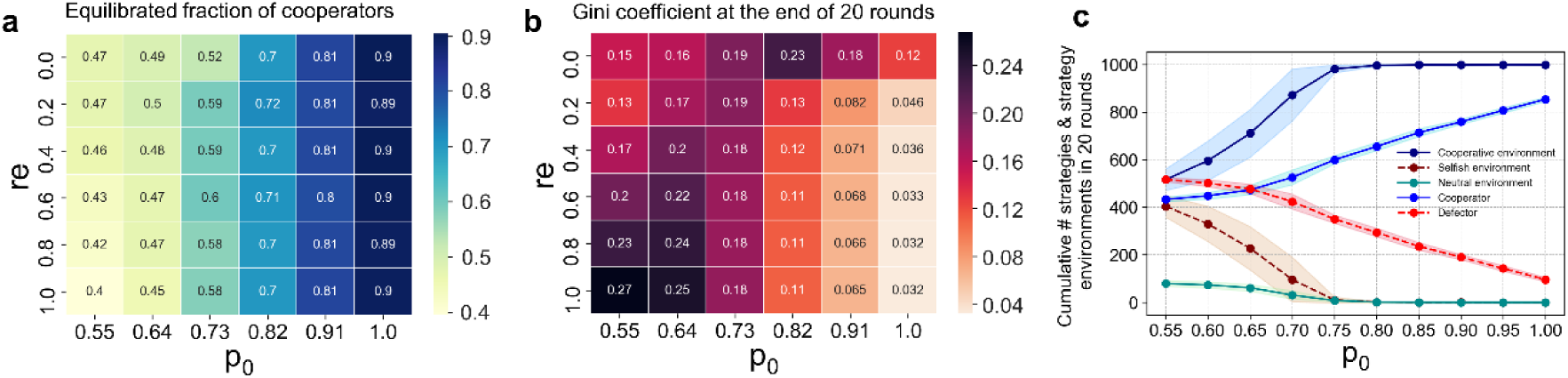
Effect of changing the influence of strategy environment on altruistic behaviour: (a) Equilibrated fraction of cooperators (b) Final Gini coefficient at the end of 20 rounds as the influence of the strategy environment (*p*_*0*_) and rewiring fraction (*re*) are varied. (c) Variation of the cumulative number of strategies and strategy environments of each type computed over all rounds of the game, with *p*_*0*_. For all panels, data points were obtained after averaging over 50 trials. The shaded regions in the last panel represent one sigma variation from the mean. Parameters: N=50, b=0.1, re=0.3,λ=0.001, r=2, initial G=0.2.

### a) Impact of the strategy environment

A greater influence of the local strategy environment in modulating cooperative behaviour increases the likelihood of cooperators to dominate over defectors. For any rewiring fraction, a higher value of p_0_ increases the propensity of rich individuals to cooperate in a cooperative environment and decreases their propensity to cooperate in a selfish environment (see Fig. 1c). The fraction of cooperators and the cumulative number of cooperative environments increase with p_0_ (Fig. 3c); this increases the number of poor as well as rich cooperators, especially at large p_0_. Since, lower wealth-inequality depends especially on the contributions of highly connected rich nodes in cooperative environments, high p_0_ (≥0.8) leads to a decrease in wealth-inequality for both static and dynamic networks, as evident from Fig. 3b. Supplementary Fig. S2 shows how the strategies, strategy environments and relative wealth of each member in the network evolves with time.

### b) Impact of remaking social ties

In a static network, poor individuals (of both C & D types) outnumber rich individuals (Fig. 4a). For moderate levels of environmental influence on decision-making (0.55≤ p_0_<0.73), increasing the network rewiring fraction (*re*) increases the number of rich defectors (RD) (Fig. 4a, also see supplementary Fig.S3(a)) while simultaneously increasing the number of cooperative environments (Fig. 4b). This results in a situation where a rich defector in a cooperative environment can exploit cooperator neighbours thereby reducing contributions to the common pool leading to lower wealth redistribution and consequently a higher level of wealth inequality (Fig. 3b).

**Figure 4:**
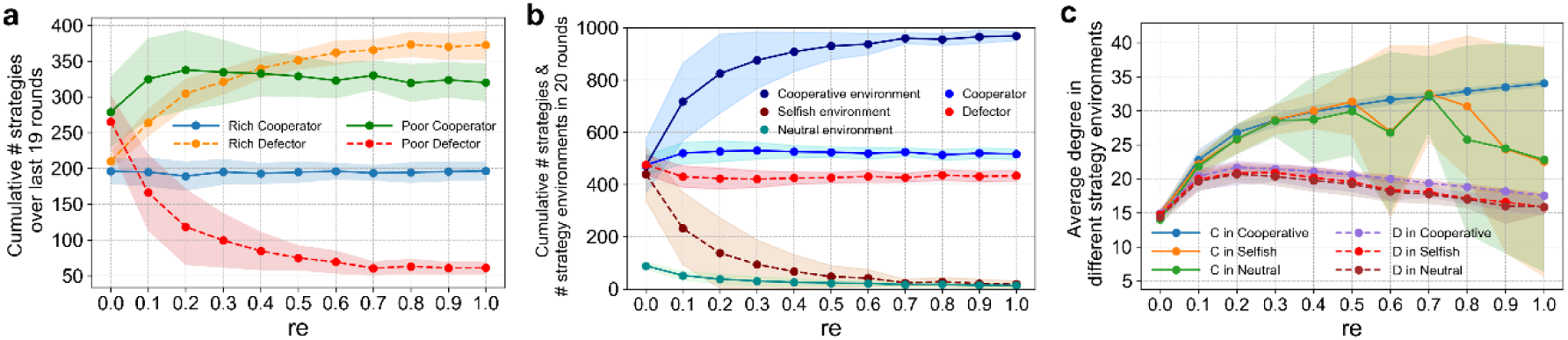
Effect of giving more individuals the option of restructuring their connections. Variation of the (a) Cumulative number of cooperators and defectors in each wealth category (b) Cumulative number of strategies (C or D) and strategy environments computed over all rounds of the game with rewiring fraction; (c) Mean degree of players in different types of strategy environments; all averaged over 50 trials. The shaded regions represent one sigma variation from the mean. With increase in rewiring rate, the degree of relatively rare C in selfish and neutral environments fluctuates wildly across different trials, which accounts for the increase in width of the shaded regions for large *re*. Parameters: N=50, b=0.1, λ=0.001, p_0_=0.7, r=2, initial G=0.2.

As the influence of the strategy environment increases, cooperators of either wealth categories dominate over defectors irrespective of the rewiring fraction (supplementary Fig.S3(c)) which increases the contribution to the common pool and facilitates efficient redistribution of wealth leading to lowering of wealth inequality. Since the average number of poor cooperators (PC) and rich cooperators (RC), who are the primary drivers for lowering wealth inequality, changes substantially only when the network changes from static to dynamic (supplementary Fig.S3) but is not affected by increase in rewiring fraction, we do not observe a significant reduction in wealth inequality with further increase in network rewiring fraction (Fig. 3b).

A highly dynamic network characterized by an increase in the rewiring of nodes leads to an increase in the number of RD primarily at the expense of poor defectors (PD) (Fig. 4a). However, these defectors have lower average degree than cooperators (Fig. 4c) and are typically embedded in a cooperative environment especially at higher p_0_ values where cooperative environments dominate over selfish ones (Fig. 4b and Fig.S4 in the electronic supplementary material). This is also reflected in the increase in number of CC links with an increase in rewiring fraction (see supplementary Fig.S5). This leads to a network where cooperators are more connected than defectors even though the local strategy environments of both strategy types are more likely to be cooperative in nature. Remarkably, the reorganization of the network topology due to the rewiring process happens in a manner that keeps the fraction of cooperators nearly constant since that only depends on the degree of influence of the local strategy environment.

### c) Effect of wealth categories on behaviour

An individual’s propensity to cooperate in the PGG can also be affected by the wealth category (rich or poor) she belongs to. Even though this dependence on relative wealth difference is weaker than the dependence on strategy environment, it reveals intriguing features of strategy update dynamics and their consequent impact on wealth inequality within the community. For example, the number of cooperators and cooperative environments remain nearly unchanged as the impact of relative wealth difference on altruistic behaviour is enhanced through increasing λ (figure not shown). However, the average number of strategy shifts (C→D, D→C) show a marked transition (Fig. 5a) from visible wealth (moderate and high λ) regime to the invisible wealth (very low λ<10^−4^) regime where the probability of cooperation is effectively independent of one’s wealth. As λ→0, in a cooperative environment, the probability of cooperation of RC increases and that of PC decreases relative to their values in the visible wealth scenario (see Eq. (2.1) & (2.2)). This makes it more likely for relatively richer individuals and less likely for relatively poorer individuals to cooperate in a cooperative environment. Since cooperative environments dominate, the former amounts to an increase in the number of RC and a decrease in the number of RD (Fig. 5b) facilitated through increasing D to C transitions. In contrast, the latter amounts to an increase in the number of PD at the expense of PC (Fig. 5b) brought about by an increase in C to D transitions. These features also explain why the final Gini increases significantly (Fig. 5c) when the visibility of wealth makes altruistic behaviour dependent on wealth categories. In such a scenario, a decrease in RC accompanied by an increase in PC and RD, even though the number of cooperative environments remain roughly constant, leads to poorer cooperators giving up some of their wealth even as a larger fraction of richer individuals increase their wealth through defection. Moreover, as the decrease in C to D and D to C transition with increasing wealth visibility indicates, RD’s and PC’s are more likely to retain their strategies over the course of the game. All these factors suppress redistribution of wealth to relatively poor individuals while favoring accumulation of wealth by richer individuals, leading to higher levels of wealth inequality. These results are consistent with the outcome of behavioural experiments which show that wealth visibility is detrimental to a reduction in wealth inequality[36].

**Figure 5:**
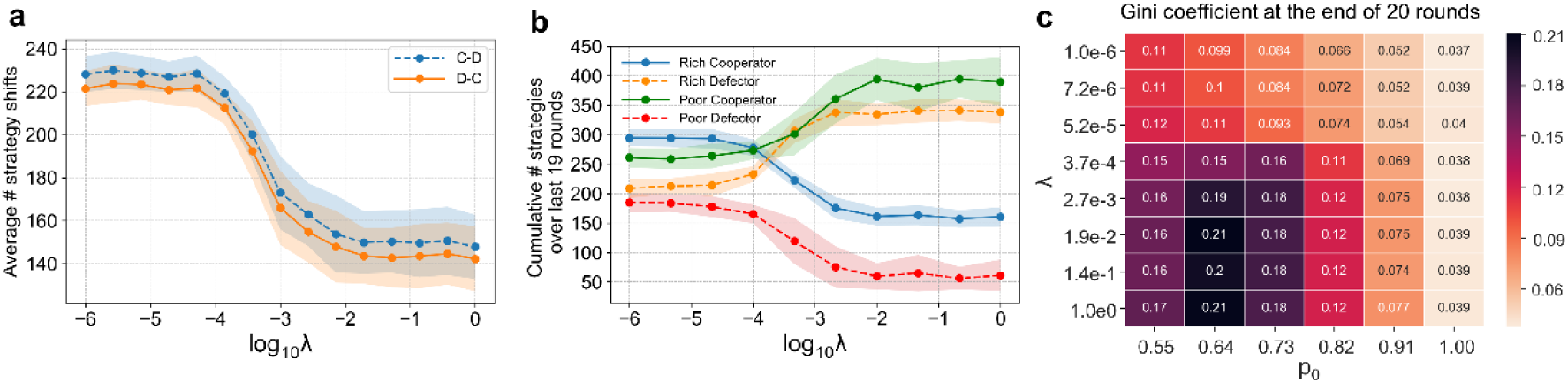
Impact of relative wealth difference on cooperative decision-making. Variation of the (a) cumulative number of C→D and D→C strategy shifts over the entire duration of the game (b) cumulative number of cooperators and defectors in each wealth category computed over all rounds of the game, with λ. (c) Heat map showing how the final wealth-inequality level at the end of 20 rounds depends on the influence of the strategy environment (p_0_) and relative wealth difference (l). All data points were obtained by averaging over 50 trials. The shaded regions in the first two panels represent one sigma variation from the mean. Parameters: N=50, b=0.1, p_0_=0.7, r_e_=0.3, r=2, initial G=0.2.

## 4. Discussion and Conclusions

Our inclinations to be altruistic are often strongly influenced by the behaviour of the people we associate with as well as our wealth relative to that of our acquaintances. Our individual response in such situations can shape collective outcomes leading to profound consequences for our own prosperity as well as the viability of sustaining public goods that rely on our contributions. Our strategy environment and wealth-dependent decision heuristic ensures that altruists are never eliminated from the population. The key aspect of our decision heuristic that ensures sustenance of cooperation is the anti-correlated nature of the propensities to cooperate in cooperative and in selfish environments. This ensures that even under unfavourable conditions characterized by relatively low probabilities of cooperation of poor individuals in either cooperative or selfish environments and of rich individuals in cooperative environments, cooperators are always sustained in the population although their mean fraction decreases slightly. When more people get the option of rewiring their social ties to punish selfish interacting partners, the change in wealth inequality levels depend on the influence of the local strategy environment on decisions to contribute in a PGG. If a cooperative strategy environment induces high propensities for cooperation from individuals regardless of their relative wealth difference, reorganization of social ties (irrespective of the rewiring fraction) can further aid in reducing wealth inequality. However, as the impact of the strategy environment on altruistic behaviour reduces, reorganization of social ties increases wealth inequality primarily because it allows RD embedded in cooperative environments to more frequently exploit their altruistic neighbours.

One way to assess the effectiveness of our decision heuristic is to examine the extent to which it can explain the results of experiments involving human subjects playing PGG on networks with the option of restructuring their social ties over time. In this context, the work of Nishi et al.[36], is particularly relevant since their experiments revealed that the visibility of wealth leads to reduced cooperation fraction and greater wealth inequality levels. Our decision heuristic is able to explain the results of those experiments for both visible and invisible wealth scenarios (see supplementary Fig.S6a-c) and thereby give us the confidence to predict that humans are indeed influenced by their strategy environment and relative wealth of the neighbourhood while updating their decisions. Despite being valid for larger population sizes, our decision heuristic is likely to be useful in populations where members possess a relatively smaller number of connected neighbours who can *influence* their decisions. This would make it possible for each person to get information about the strategy environment and wealth of such neighbours. For very large network neighbourhoods, such information would be more difficult to acquire, process and utilize due to cognitive limitations. The population sizes we have chosen are therefore of the order typically sampled in behavioural experiments[36,37,39,40]. Such a choice seems reasonable since a lot of social interactions occur within relatively small communities where individual behaviour is shaped by a few influential interacting partners.

Since our decision heuristic imposes distinctly different propensities to cooperate for rich and poor members of the community in different strategy environments, we also compared the outcomes of alternative decision heuristics to examine the impacts of each on cooperation levels and wealth inequality. One such alternative makes richer and poorer individuals more likely to cooperate in cooperative and selfish environments respectively. Then, populations with higher initial Gini saturate to lower and unrealistic final wealth inequality levels not supported by experiments[36] (see supplementary Fig.S6d), than those with lower initial Gini.

To check the robustness of our results, we have verified the effect of variation in other parameters like synergy factor, likelihoods of making, breaking and retaining links and population size. An increase in synergy factor increases average wealth and reduces wealth inequality, without affecting the fraction of cooperators or average degree of the network, since a larger multiple of the altruist’s contribution is redistributed to each connected neighbour. Our results are also mostly robust to changes in network restructuring probabilities (see supplementary Fig.S7). If however, rigid rules for network restructuring are imposed, with existing links retained and new links made only if both connecting partners are cooperators, a more dynamic network (larger *re*) is counter-productive for sustaining cooperation and even maintaining social cohesion due to breakdown of social ties (see supplementary Fig.S8, S9).

Even though we have not explicitly included punishing strategies[44], it is indirectly accounted for through the rewiring of social ties. Such a process can be thought of as a cost-free mechanism of penalizing selfish behaviour; whose effect is manifest through the lower average degree of defectors in comparison to the cooperators. It will be interesting to see how explicit incorporation of punishing strategies in our decision-making framework affects the strategy distribution and wealth inequality levels in the population. Introducing heterogeneity in contributions for the upkeep of the public good, which is akin to imposing wealth-dependent tax brackets, may also constitute a fairer mechanism for encouraging altruistic behaviour.

Altruistic behaviour in humans can depend on a variety of motivating factors such as empathy, personal values and aspirations, social environment and cultural norms. The relative importance of these diverse factors can not only vary from one person to another but also depend on the nature of the social dilemma encountered. While no single decision rule can fully capture the behavioural complexities of individuals, by studying emergent patterns in human behaviour in different situations, it might be possible to discover general principles of decision-making that can then be encapsulated in a heuristic. The decision-making heuristic proposed and explored here provides a simple, yet powerful framework for understanding how the co-evolution of individual choices and social network ties can work in tandem to sustain cooperation levels and consequently affect wealth inequality in the population.

## Author Contributions

SP, PV, SS conceived and designed the project. SKR contributed to initial exploration and code development. SP and PV developed the model with inputs from SS. SP developed code in Python and carried out the simulations. PV developed code in Matlab and validated the simulation results. SP, PV and SS analysed the simulation results. SS wrote the paper with inputs from SP, PV, SKR.

## Acknowledgements

We thank Anuran Pal for helpful discussions.

## Notes

### Competing Interest Statement

The authors have declared no competing interest.

### Summary of Updates

The introduction and discussion sections have been significantly modified to improve readability.

