## supplementary material for "How strategy environment and wealth shape altruistic behaviour: Cooperation rules affecting wealth distribution in dynamic networks"

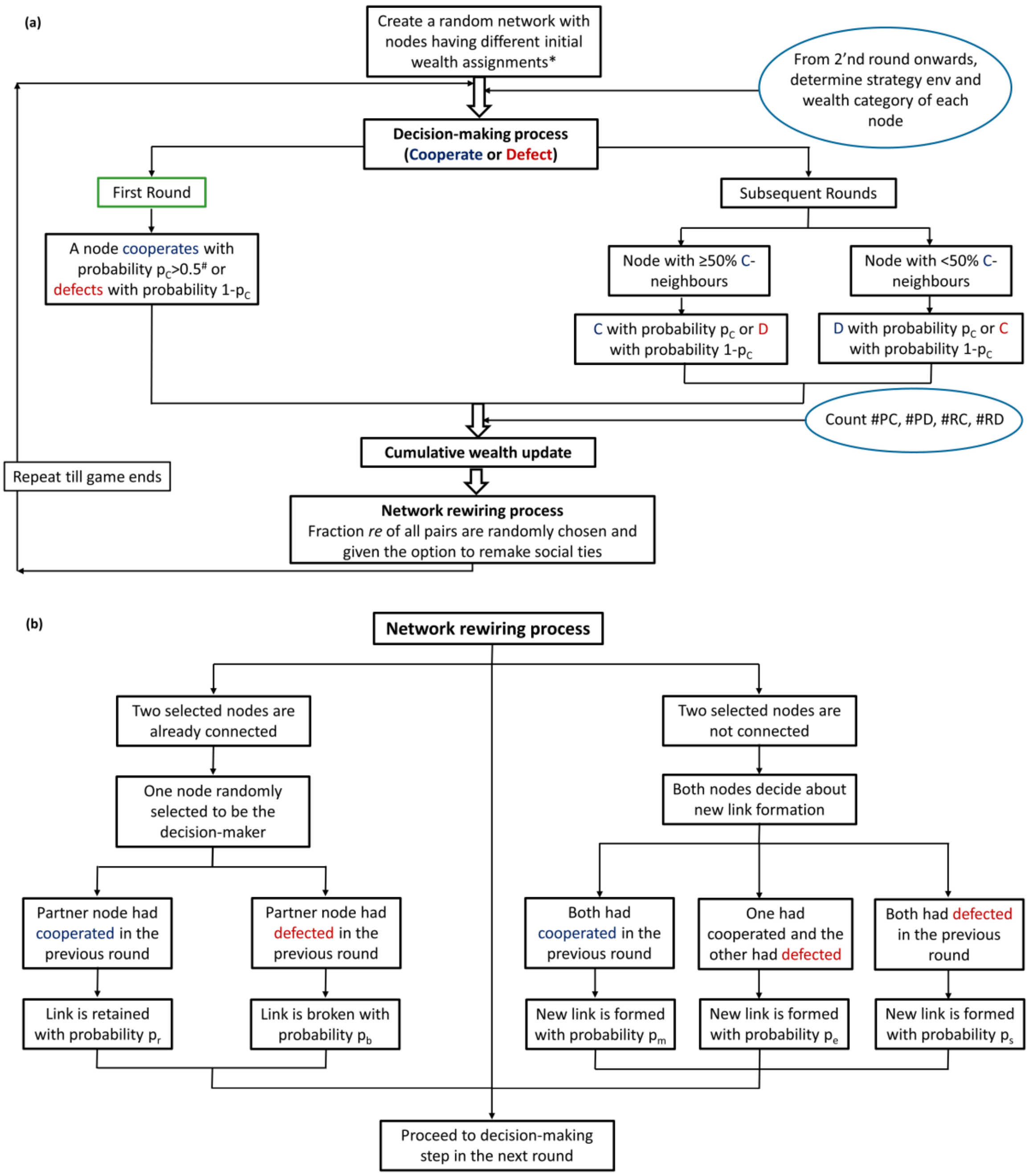


**Figure S1:** (a) An algorithmic view of the model. (b) Details of the network restructuring process. PC: Poor Cooperators, PD: Poor Defectors, RC: Rich Cooperators, RD: Rich Defectors.

*Initial wealth assignment to each individual depends on the initial Gini. Following Nishi et al.[1], when (a) initial Gini=0, each individual is given a fixed amount of money = 500 units (initial endowment). (b) initial Gini=0.2, each member of the community is assigned one of the two wealth categories (rich or poor) with a probability f=0.5 with rich and poor individuals getting 700 and 300 units respectively. (c) initial Gini=0.4, each member of the community is assigned to be rich with a probability f=0.3 with rich and poor individuals getting 1150 and 200 units respectively.

#For all simulations, the value of $p_C$ in the first round was chosen to be between 0.6 and 0.7. Change in this value does not affect the final conclusions.

*Metrics used to analyze the evolutionary dynamics*: The evolution of altruistic behaviour and wealth inequality level in the population was analyzed by varying the effect of strategy environment, wealth and rewiring fraction, the three crucial parameters in the model. The key metrics tracked are defined as follows.

The cumulative number of strategies (C or D) and strategy environments (cooperative, selfish or neutral) is given by $\#S_a = \frac{1}{N_T}\sum_{i=1}^{N_T}\left[\sum_{j=1}^{M}\#S_a(j)\right]$ and $\#S_e = \frac{1}{N_T}\sum_{i=1}^{N_T}\left[\sum_{j=1}^{M}\#S_e(j)\right]$ respectively; where $\#S_a(j)$ and $\#S_e(j)$ give the number of strategies (C or D) and strategy environments (cooperative, selfish or neutral) respectively, in the j'th round.

The average degree of the network in the j'th round: $<k(j)>= \frac{1}{N_T}\sum_{i=1}^{N_T}\frac{\sum_{n=1}^{N}k_n^i(j)}{N}$

The average degree of a C or D in different strategy environments is given by

$<k_X^Y>= \frac{1}{N_T}\sum_{i=1}^{N_T}\left[\frac{\sum_{j=1}^{M}\sum_{l=1}^{S}k_{X_l}^Y(j)}{\sum_{j=1}^{M}n_X^Y(j)}\right]$ where X=C or D and Y=C- or D- or neutral-environment. $k_{X_l}^Y(j)$ is the degree of the $l$'th X in Y-environment in the $j$'th round and $S = n_X^Y(j)$ is the number of X's in Y-environments in the $j$'th round, M is the total number of rounds and $N_T$ denotes the number of trials. The average number of strategy shifts $<S_S>$ is given by $<S_s>= \frac{1}{N_T}\sum_{i=1}^{N_T}\left[\sum_{\substack{j=1\\X\neq Y}}^{M}\#(X\rightarrow Y)(j)\right]$ where #(X➔Y)($j$) gives the number of strategy shifts (C➔D or D➔C) in the $j$'th round.

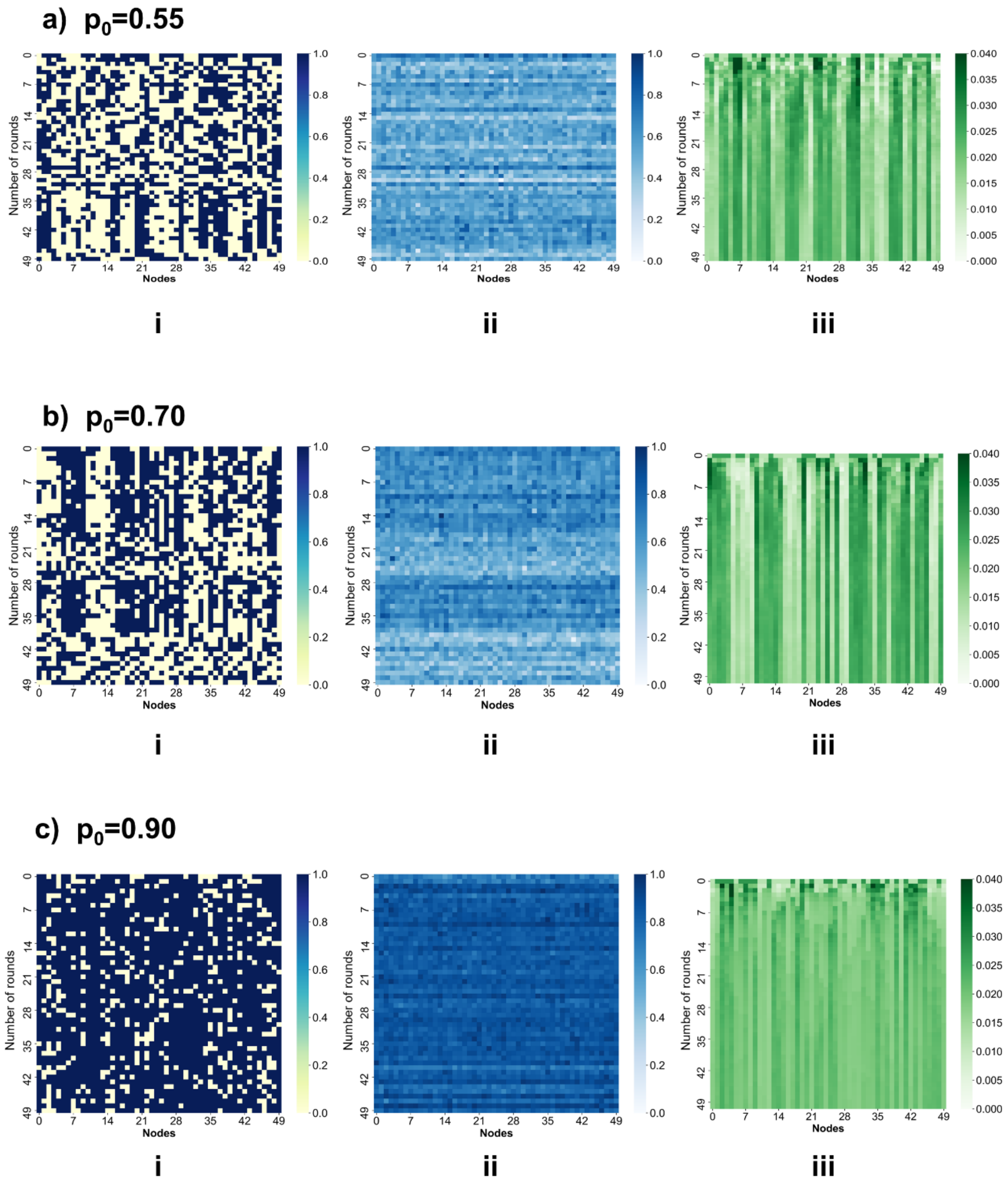


**Figure S2:** Time evolution of (i) individual strategies where dark blue signifies cooperators and pale yellow indicates defectors (ii) individual strategy environments with darker shades of blue indicating a larger fraction of cooperator neighbours. (iii) individual wealth as a fraction of the total wealth with darker shades of green denoting richer individuals; for a single trial, in a *dynamic* network, for different values of the environment factor $p_0$. Other parameters are N=50, $\lambda$=0.001, b=0.1, re = 0.3, r=2, initial Gini = 0.2

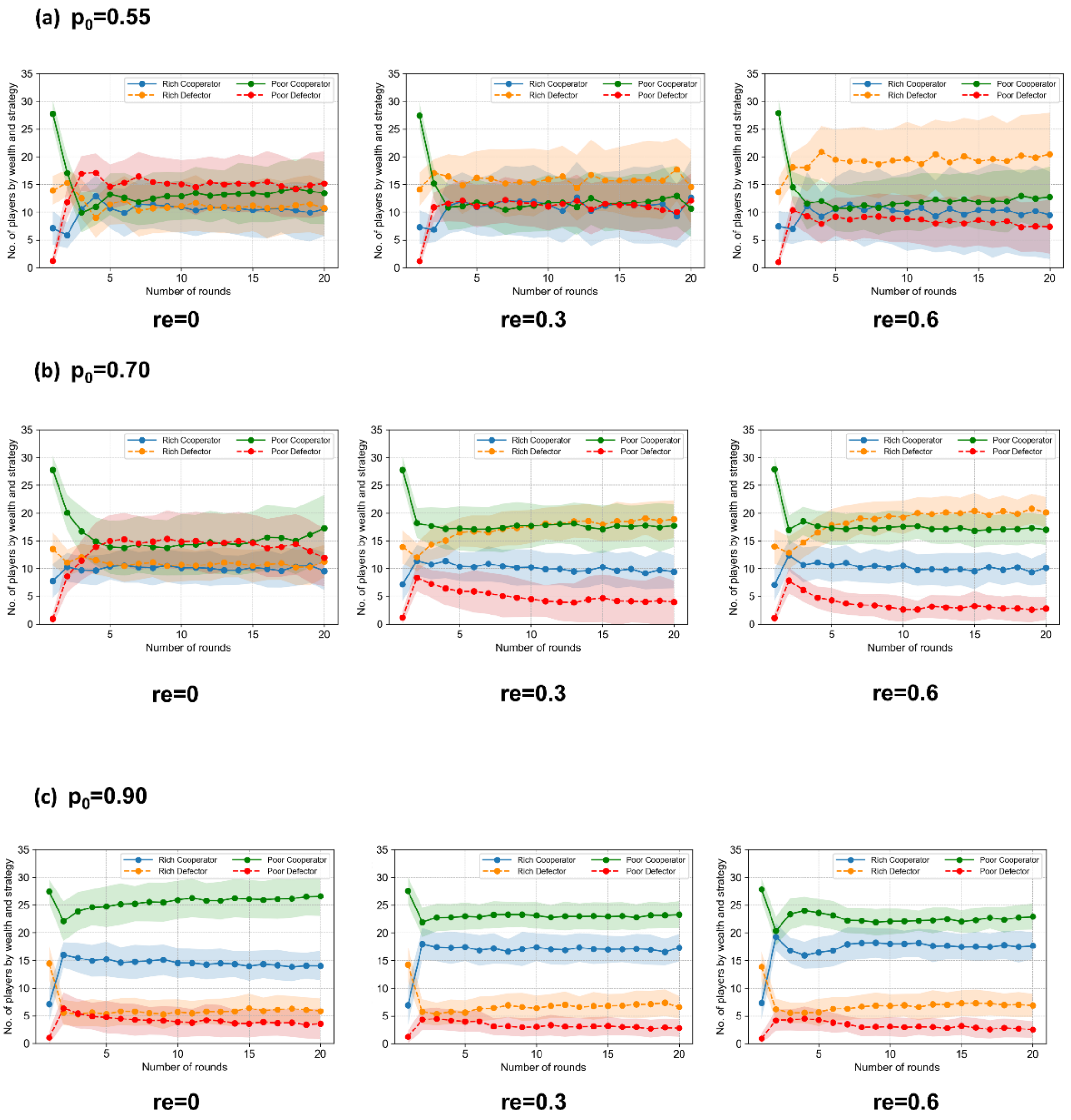


**Figure S3:** Time evolution of cooperators and defectors belonging to different wealth categories, each data point is obtained by averaging over 100 trials. The shaded regions represent one sigma variation from the mean. Results for 3 different rewiring probabilities are shown for each value of the environment factor (a) $p_0$ = 0.55, (b) $p_0$ = 0.7, (c) $p_0$ = 0.9. Other parameters are N=50, $\lambda$=0.001, b=0.1, r=2, initial Gini = 0.2.

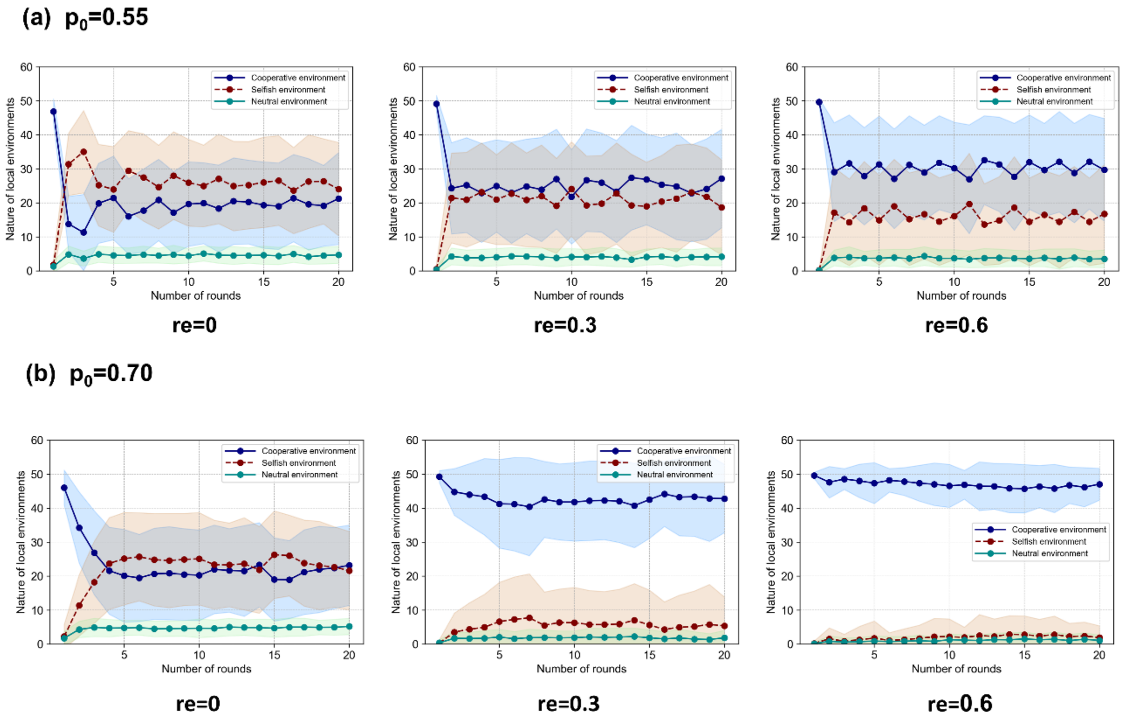


**Figure S4:** Time evolution of the number of strategy environments, each data point is obtained by averaging over 100 trials. A cooperative, selfish or neutral environment is defined by fraction of cooperator neighbours of the focal player being >0.5, <0.5, =0.5 respectively. The shaded regions represent one sigma variation from the mean. Results for 3 different rewiring probabilities are shown for each value of the environment factor (a) $p_0$ = 0.55, (b) $p_0$ = 0.7, Other parameters are N=50, $\lambda$=0.001, b=0.1, r=2, initial Gini = 0.2

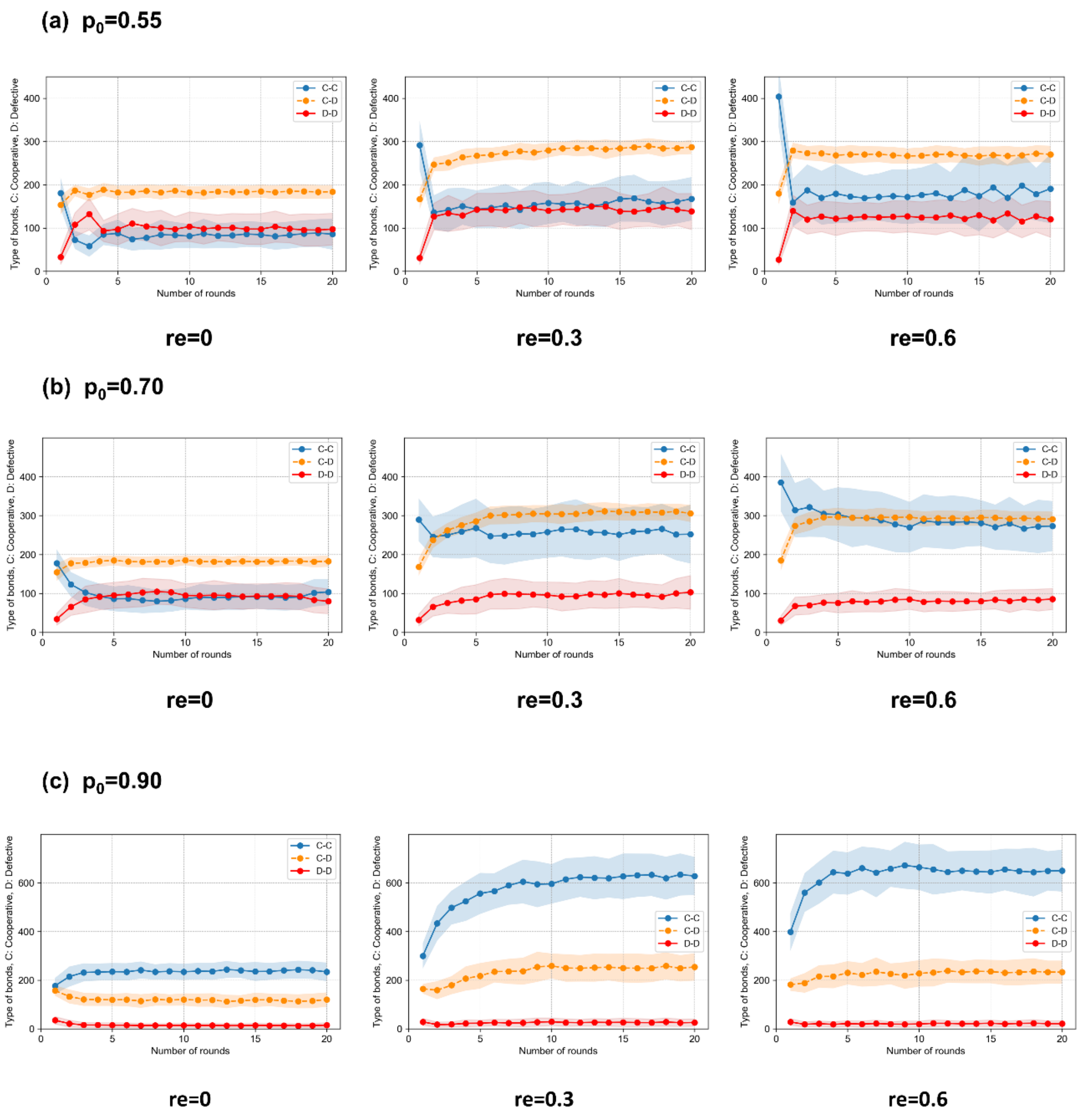


**Figure S5:** Time evolution of number of CC, CD, DD links, each data point is obtained by averaging over 100 trials. The shaded region represents one sigma variation from the mean. Results for 3 different rewiring probabilities are shown for each value of the environment factor (a) $p_0$ = 0.55, (b) $p_0$ = 0.7, (c) $p_0$ = 0.9. Other parameters are N=50, $\lambda$=0.001, b=0.1, r=2, initial Gini = 0.2

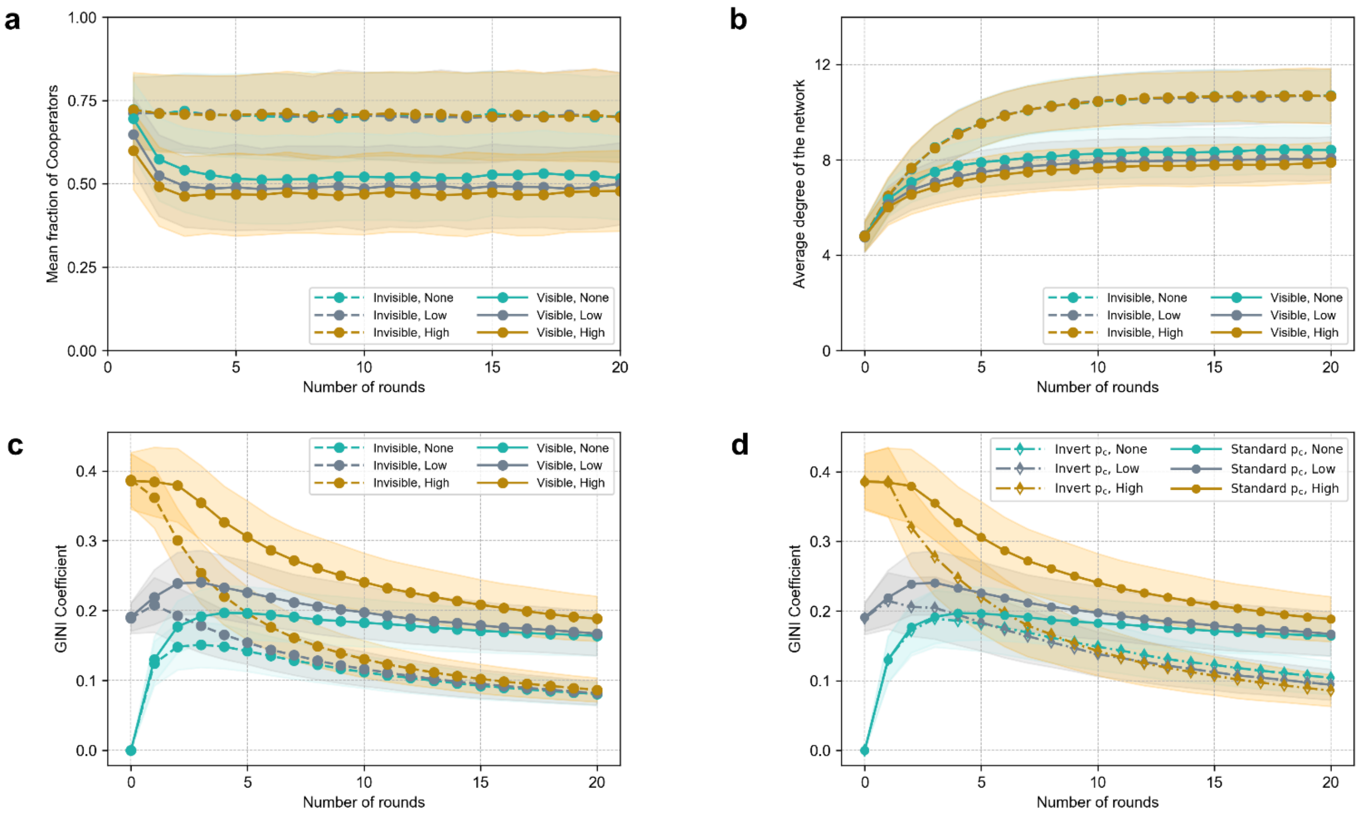


**Figure S6:** Time evolution (averaged over 1000 trials) of the (a) mean fraction of cooperators (b) average degree (c) wealth-inequality measured by the Gini coefficient for our decision heuristic as described by Eq. 3&4. The dotted lines correspond to the invisible wealth scenario characterized by $\lambda$=0 and the solid line corresponds to visible wealth scenario where the likelihood of cooperation of an individual depends on the wealth category she belongs to. The shaded region represents one sigma variation from the mean. We use a small network size of N=17 to enable direct comparison with the experimental results reported in Nishi et al.[1] (d) Comparison between our standard decision heuristic shown in *solid* lines and the inverted decision heuristic where Eq. 3 & 4 gives the probability of cooperation of the focal player in selfish and cooperative environments respectively, shown in *dash-dotted* lines. Both sets of lines correspond to the visible wealth scenario with $\lambda$=0.001. Other parameters are: r = 2, re = 0.3, b=0.1, initial Gini of 0 (None), 0.2 (Low) and 0.4 (High) for which p0=0.69, 0.65 and 0.6 respectively.

In the *inverted* decision heuristic, richer individuals are more likely to cooperate in a cooperative environment and poorer individuals are more likely to cooperate in selfish environments. In this case, the probability of cooperation ( $p_C$ ) for a focal player is given by

$p_C = p_0 - b + (1 - p_0)\tanh(\lambda \Delta w)$; when the focal player is in a selfish environment and

$p_C = (1 - p_0 + b) - (1 - p_0)\tanh(\lambda \Delta w)$; when the focal player is in a cooperative environment.

As before, $\Delta w = E_{LC}(w) - w_F$ is the relative wealth difference; $E_{LC}(w)$ is the average cumulative wealth of neighbours of the focal player; $w_F$ is the focal player's cumulative wealth accumulated over past rounds. The outcome of this inverted decision heuristic on wealth inequality is represented by the dash-dotted lines in the above figure. The final Gini coefficient are not consistent with the results of experiments described in Nishi et al.[1].

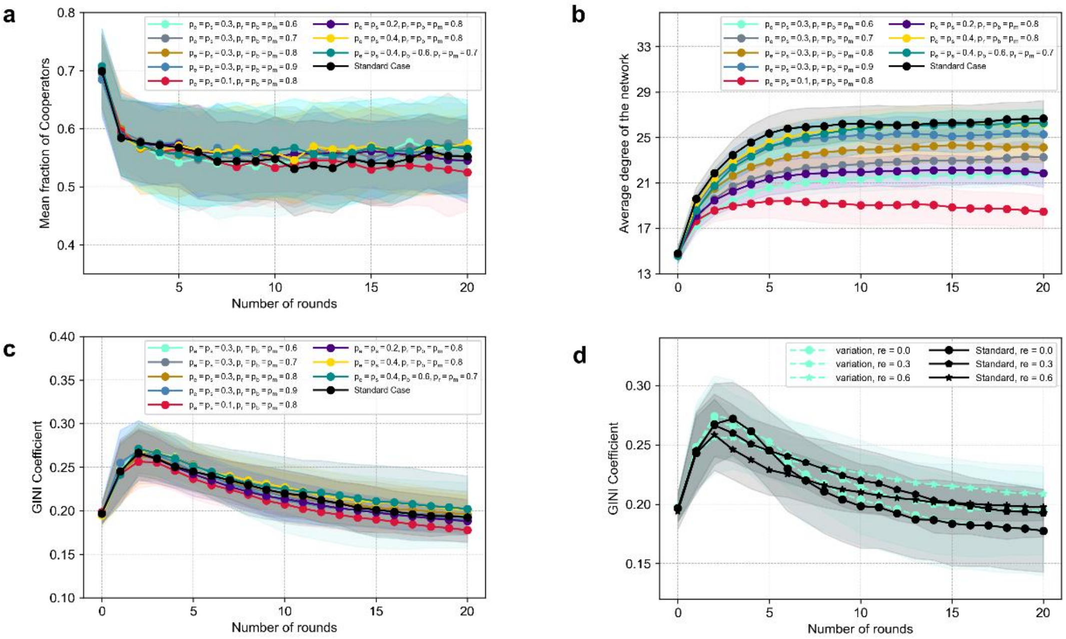


**Figure S7: Effect of varying network restructuring probabilities.** Time evolution (averaged over 100 trials) of the (a) mean fraction of cooperators (b) average degree and (c) Gini coefficient for fixed rewiring fraction (re=0.3) but different values of network restructuring probabilities. (d) Gini coefficient for different rewiring fractions and two distinct sets of network restructuring probabilities. The black curve corresponds to the standard restructuring probabilities ($p_r$=0.87, $p_m$=0.93, $p_b$=0.7, $p_e$=0.3, $p_s$=0.2) used to generate all other plots in the manuscript. The light green curves in panel d correspond to first set of network restructuring probabilities shown in panels a, b &c. Other parameters are: N=50, r=2, b=0.1, $\lambda$=0.001, p0=0.7, initial Gini = 0.2.

Different rewiring probabilities used in the above example does not affect the qualitative trends in cooperator fraction, final Gini coefficient and average degree. Even the slight changes in the final values of the first two quantities are within 1-sigma margins of error. The changes in the equilibrium value of the average degree of the network are somewhat more pronounced with the average degree decreasing systematically as network restructuring probabilities approach the extreme values characteristic of rigid rewiring rules (compare red line in S7(b) with gray line in Figure S8(c)).

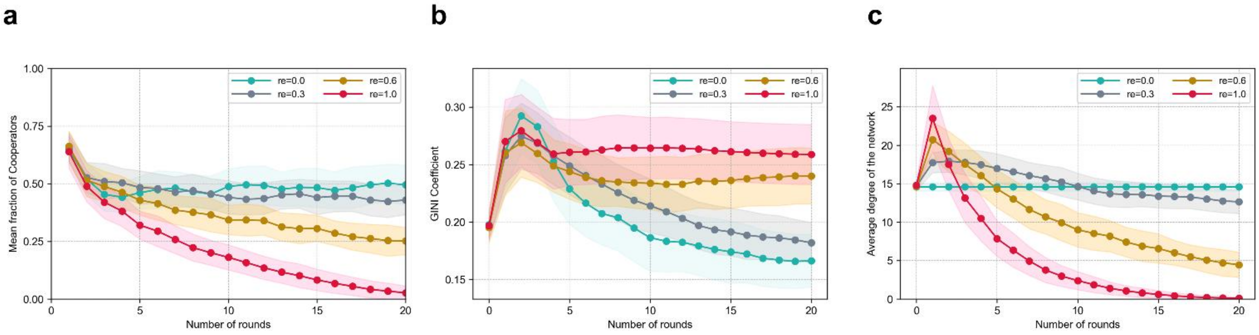


**Figure S8:** Time evolution (averaged over 100 trials) of the (a) mean fraction of cooperators (b) Gini coefficient and (c) average degree for static and dynamic networks when deterministic rules are imposed for retaining, making and breaking links. The network restructuring probabilities are $p_r$=$p_m$=$p_b$=1, $p_e$=$p_s$=0. Other parameters are: N=50, r = 2, b=0.1, $\lambda$=0.001, p0=0.65, initial Gini=0.2.

When existing links are retained and new links are made only if the opponent is a cooperator, increase in rewiring fraction (*re*) has dramatic effects on fraction of cooperators, Gini coefficient and average degree as evident from the above plot. These trends can be understood by observing how the number of RD, cooperative environments and CC, CD links are affected by large increase in rewiring fraction (see Figure S9). The number of RD increases at first as the rewiring fraction increases, since RD have a higher propensity to defect in a cooperative environment. Since cooperative strategy environments dominate, this allows rich defectors to increase their wealth by exploiting altruistic neighbours, thereby increasing wealth inequality in the population. At the same time, rigid rewiring probabilities lead to altruists as well as selfish individuals breaking their links with other defectors (when given the option to do so; which becomes more likely when *re* is high) resulting in a less connected network with smaller number of CD and DD links. The number of CC links also decreases because of the overall decrease in the number of both poor and rich altruists. This combination of factors leads to a population with higher levels of wealth inequality and a network with large number of defectors with fewer connected neighbours on an average. These effects get amplified as the rewiring fraction increases further, eventually leading to complete breakdown of social ties in the limit *re* ➔ 1.

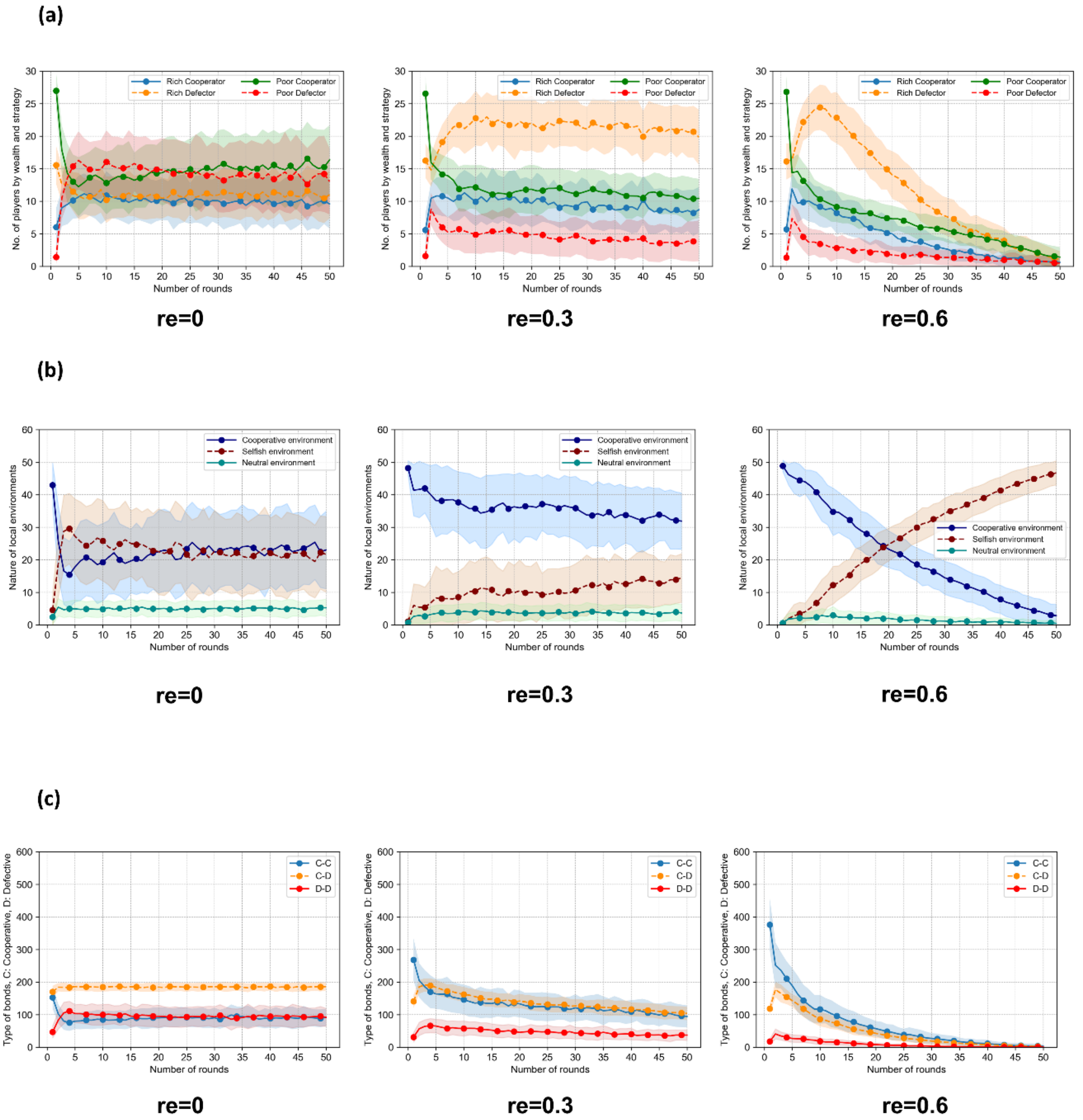


**Figure S9:** Time evolution (averaged over 100 trials) of the (a) number of players distinguished by their strategy and wealth category (b) number of strategy environments of each type (c) number of CC, CD and DD links; when deterministic rules are imposed for retaining, making and breaking links i.e. the network restructuring probabilities are $p_r$=$p_m$=$p_b$=1, $p_e$=$p_s$=0. Other parameters are: N=50, r = 2, b=0.1, $\lambda$=0.001, p0=0.65, initial Gini=0.2.

**Supplementary Video1:** The movie shows the evolution of strategies and wealth on a 12-node dynamical social network. Other parameters are: $p_0$ = 0.6, $\lambda$=0.001, $r_e$ = 0.3, b=0.1, r=2, initial Gini = 0.

### Supplementary material References

1. Nishi A, Shirado H, Rand DG, Christakis NA. 2015 Inequality and visibility of wealth in experimental social networks. *Nature* **526**, 426–429. (doi:10.1038/nature15392)
